# Identifying Developmental Changes in Functional Brain Connectivity Associated with Cognitive Functioning in Children and Adolescents with ADHD

**DOI:** 10.1101/2023.12.20.572617

**Authors:** B Pho, RA Stevenson, Y Mohzenszadeh, B Stojanoski

**Affiliations:** Program in Neuroscience, University of Western Ontario, London, ON; Brain and Mind Institute, University of Western Ontario, London, ON; Department of Psychology, University of Western Ontario, London, ON; Western Institute for Neuroscience, University of Western Ontario, London, ON; Department of Computer Science, Western University, London ON, N6A 5B7, Canada; Vector Institute for Artificial Intelligence, Toronto, Ontario, Canada; Faculty of Social Science and Humanities, Ontario Tech University, Oshawa, ON

## Abstract

Children and adolescents diagnosed with Attention Deficit Hyperactivity Disorder (ADHD) often show deficits in various measures of higher-level cognition, such as, memory and executive functioning. Poorer high-level cognitive functioning in children with ADDH has been associated with differences in functional connectivity across the brain, including the frontoparietal network. However, little is known about the developmental changes to cortical functional connectivity profiles associated with higher-order cognitive abilities in this cohort. To characterize changes in the functional brain connectivity profiles related to higher-order cognitive functioning, we analyzed a large dataset(n=479) from the publicly available Healthy Brain Network which included fMRI data collected while children and adolescents between the ages of 6 and 16 watched a short movie-clip. The cohort was divided into two groups, neurotypical youth (n=106), and children and adolescents with ADHD (n=373). We applied machine learning models to functional connectivity profiles generated from the fMRI data to identify patterns of network connectivity that differentially predict cognitive abilities in our cohort of interest. We found, using out-of-sample cross validation, models using functional connectivity profiles in response to movie-watching successfully predicted IQ, visual spatial, verbal comprehension, and fluid reasoning in children ages 6 to 11, but not in adolescents with ADHD. The models identified connections with the default mode, memory retrieval, and dorsal attention networks as driving prediction during early and middle childhood, but connections with the somatomotor, cingulo-opercular, and frontoparietal networks were more important in middle childhood. This work demonstrated that computational models applied to neuroimaging data in response to naturalistic stimuli can identify distinct neural mechanisms associated with cognitive abilities at different developmental stages in children and adolescents with ADHD.

## Introduction

Attention Deficit Hyperactivity Disorder (ADHD) is the most common neurodevelopment disorder among children and adolescents, affecting an estimated 4.8% of all Canadian children up to 19 years of age ^1^. One reason ADHD is commonly diagnosed in school-aged children is because the symptoms linked to ADHD are most salient in the classroom ^2^. For instance, ADHD is best characterized by a persistent pattern of inattention (inability to maintain focus), impulsivity (acting on instinct without thinking), and/or hyperactivity (excessive restlessness and movement) that can interfere with not only completing school-based tasks, but extends to daily functioning ^3–6^

One of the most common aspects of ADHD is a deficit in processing speed ^7–12^, and executive functioning, which is comprised of three components: inhibitory control, cognitive flexibility, and working memory ^11,13,14^. Indeed, recent studies have found converging evidence that the brain-based associations of ADHD include reduced activity in prefrontal cortex, basal ganglia, cerebellum, and parieto-temporal regions, all of which have been shown to support multiple cognitive processes such as cognitive control, working memory, and attention ^15–17^.

Although there is a large literature examining the cognitive abilities in individuals with ADHD, many of those studies have focused on adults or single-age cohorts of children, without considering development. Consequently, less is known about the neural mechanisms associated with cognitive development from childhood to adolescence in individuals with ADHD.

Advancements in applying machine learning to large neuroimaging datasets has proven to be a valuable tool to understand the relationship between cognition and the underlying neural mechanisms^18–24^. For example, machine learning (i.e., Ridge regression) has been successfully applied to resting-state functional connectivity networks to predict fluid and crystalized intelligence in healthy young adults ^25^. A similar approach was also used to predict links between neural activity in the default mode network and three task control networks (frontoparietal, salience, and dorsal attention) and higher-order cognitive functions, such as, general ability, speed/flexibility, and learning/memory in younger participants^26^. Although much of this work on applying machine learning to link neural activity with cognitive ability has relied on resting-state data, movie-watching fMRI has been shown to improve functional connectivity-based prediction of behavior compared to resting-state^27–29^. The few studies that have used movie-watching data were able to successfully predict cognitive abilities in neurotypical adult^29^ and child populations^30^. The advantage of using movie-watching data is likely due to reduced motion, increased engagement, but perhaps most importantly, movie-watching requires the integration of various cognitive systems to follow the complexities of the plot. Moreover, individuals often have a unique interpretation of the movie, resulting in enhanced individual signals and therefore richer brain dynamics can be captured by predictive models^31–33^.

In the current study, we combined movie-watching fMRI and machine learning to identify different patterns of functional network connectivity that best predict cognitive ability in a large cohort of children and adolescents with ADHD. We predicted that not only are there specific neural mechanisms associated with different aspects of higher-level cognition in children and adolescents with ADHD, but those mechanisms change developmentally and are unique to different age groups. To explore changes in the neural mechanisms associated with cognitive functioning across time, participants were divided into three age bins and neural activity was modeled (in response to a movie) to predict the same set of cognitive abilities for each age bin. By splitting participants into three age bins, the models would either 1) predict the same set of cognitive abilities for all three age bins, suggesting a similar functional connectivity profile across development or 2) predict a different set of cognitive abilities for each age bin, suggesting the model captured a functional connectivity profile unique to age cohorts. We compared models using out-of-sample cross-validation (model trained on one age bin to predict the same cognitive ability in a different age bin) to determine the degree to which similar neural properties were associated with cognition across age bins. We analyzed shared functional connectivity profiles by calculating a difference score between the models (feature weights) trained on each age bin, revealing connections that changed the most or the least across age bins.

## Methods and Materials

### Participants

We obtained data from the Healthy Brain Network (HBN) biobank (releases 1 to 8) as part of the Child Mind Institute^34^. The Chesapeake Institutional Review Board approved the study, and details on the HBN biobank can be found at: http://fcon_1000.projects.nitrc.org/indi/cmi_healthy_brain_network/

We included a sample of 479 data sets from children and adolescents between the ages 6 to 16 in the final analysis. The data sets consisted of a T1-weighted, and functional MRI scan, along with phenotypic data. We excluded participants with lower-quality data, based on visual inspection of the T1 images and functional connectivity matrices, along with those who full-scale intelligent quotient scores under 70 (Supplement).

Phenotypic data included age, sex, clinical diagnosis, and six cognitive measures from the Wechsler Intelligence Scale (Table 1). Clinical diagnoses were provided by up to ten licensed clinicians after interviews with the parents and child^34^ which we used to group participants into the ADHD (at least one diagnosis of “ADHD”) or NT (no clinical diagnoses) group. In addition to a single ADHD group (n=373), we divided participants with ADHD into three age bins: early childhood (Bin 1: ages 6-8, n=114), middle childhood (Bin 2: ages 9-11, n=147), and adolescence (Bin 3: ages 12-16, n=112). Due to the smaller sample size (n=106), we did not divide the Neurotypical (NT) group into discrete age bins.

**Table 1:** Participant demographics. For each group and measure, the mean, standard deviation (in brackets), and range are provided.

|  | **Group** |  |  |  |  |
| --- | --- | --- | --- | --- | --- |
|  | **ADHD** |  |  |  | **TD** |
| **Measure** | **All** | **Bin 1** | **Bin 2** | **Bin 3** | **All** |
| **N** | 373 | 114 | 147 | 112 | 106 |
| **Age** | 10.57 (2.53)<br>6.03 to 15.98 | 7.73 (0.76)<br>6.03 to 8.98 | 10.34 (0.88)<br>9.04 to 11.96 | 13.75 (1.13)<br>12.03 to 15.98 | 10.12 (2.78)<br>6.05 to 16.50 |
| **Sex (M/F)** | 274/99 | 77/37 | 118/29 | 79/33 | 62/44 |
| **WISC FSIQ** | 100.13 (16.31)<br>70.00 to 147.00 | 104.13 (15.47)<br>73.00 to 138.00 | 98.56 (15.90)<br>70.00 to 141.00 | 98.11 (16.94)<br>71.00 to 147.00 | 108.58 (14.20)<br>76.00 to 145.00 |
| **WISC VSI** | 102.32 (17.12)<br>57.00 to 155.00 | 106.38 (16.64)<br>67.00 to 147.00 | 100.73 (16.69)<br>64.00 to 144.00 | 100.28 (17.46)<br>57.00 to 155.00 | 105.21 (15.01)<br>64.00 to 141.00 |
| **WISC VCI** | 105.04 (15.87)<br>70.00 to 155.00 | 108.53 (16.06)<br>70.00 to 146.00 | 104.44 (15.34)<br>70.00 to 155.00 | 102.29 (15.71)<br>70.00 to 142.00 | 110.58 (13.93)<br>78.00 to 155.00 |
| **WISC FRI** | 101.33 (16.19)<br>67.00 to 144.00 | 104.58 (15.40)<br>67.00 to 140.00 | 99.14 (15.75)<br>67.00 to 134.00 | 100.90 (16.98)<br>67.00 to 144.00 | 107.42 (15.11)<br>69.00 to 155.00 |
| **WISC WMI** | 98.54 (15.18)<br>62.00 to 138.00 | 99.53 (14.83)<br>67.00 to 138.00 | 97.21 (14.04)<br>65.00 to 138.00 | 99.29 (16.77)<br>62.00 to 135.00 | 103.92 (14.29)<br>72.00 to 135.00 |
| **WISC PSI** | 93.65 (15.35)<br>53.00 to | 97.33 (15.29)<br>56.00 to | 93.12 (13.94)<br>56.00 to | 90.59 (16.38)<br>53.00 to | 106.55 (15.80)<br>66.00 to |
|  | 148.00 | 148.00 | 123.00 | 132.00 | 155.00 |

### (f)MRI acquisition and preprocessing

T1-weighted anatomical and functional MR images were acquired while participants watched a ten-minute clip from the movie ‘Despicable Me’^34^. Neuroimaging data were preprocessed and analyzed using the Automatic Analysis (AA) toolbox^35^, SPM 8, and in-house MATLAB scripts (see Supplement for additional details).

We generated functional connectivity matrices for each participant using 264 regions-of-interest (ROI) as defined in the Power et al. (2011) atlas^36^. Individual ROIs comprised of spheres of 5 mm in radius with spatial smoothing full-width half maximum of 6 mm and z-score standardization. We correlated activity in each sphere to every other sphere, resulting in a 264 x 264 functional connectivity matrix.

### Cognitive ability

Cognitive ability was measured using the Wechsler Intelligence Scale for Children Fifth Edition^37^. The WISC-V measures a child’s intellectual ability based on five primary indices: Visual Spatial Index (VSI), Verbal Comprehension Index (VCI), Fluid Reasoning Index (FRI), Working Memory Index (WMI), and Processing Speed Index (PSI). In addition, the WISC-V also provides a Full-Scale IQ (FSIQ) score, which is derived from the five primary indices, and normalized by age (description of the cognitive measures can be found in the Supplement).

### Computational Modeling

We used two computational models to examine the relationship between functional connectivity and cognitive ability: partial least squares (PLS)^38,39^ and Ridge regression^40,41^. Ridge and PLS models are ideal for high-dimensional multicollinear data and have built-in anti-overfitting (regularization) parameters (details about optimal components and alpha values in Supplement).

We first fit standard scaler models to rescale features such that they have the properties of a standard normal distribution with a mean of zero and a variance of one. This is essential because regularized linear models assume features are centered around zero and have variance in the same scale to avoid certain features dominating because of differences in variance. To avoid data leakage between the training and testing set, we fit the standard scaler only on the training set, and it was applied to both the training and testing set.

### Model Feature Weight Analysis

After we trained the models, we analyzed the model’s feature weights using two methods. First, we assessed feature weight reliability between different computational models using the intraclass correlation coefficient^25^ (ICC). In the second method, we used feature weights trained on one subset of the dataset and applied them to predict cognition on a different subset. We based prediction accuracy on Pearson correlations representing the degree of similarity between the model’s predicted values of cognitive ability and the true values. We calculated statistical significance by comparing the observed Pearson r score relative to a null distribution of Pearson r scores generated from 500 random permutations of the dataset. We performed this out-of-sample cross-validation—referred to as cross-prediction—only on the ADHD group and evaluated it using permutation statistics (Supplement).

Model feature weights represent the weight (importance) associated with specific aspects of the functional connectivity matrix. We multiplied the feature weights by a participant’s functional connectivity matrix and used this to calculate the predicted score across all cognitive measures. We then compared this score against the participant’s actual cognitive score to update the feature weights. Thus, the feature weights represent a heat map of important functional connections for predicting cognitive ability.

To explore how cognition develops, we subtracted feature weights of models trained on early childhood (Bin 1) from models trained on middle childhood (Bin 2), with respect to cognitive ability. We used absolute values to highlight the magnitude of change between Bin 1 and Bin 2 feature weights. The absolute-value-feature-weight-differences matrix represents the network connections that change in importance between Bins 1 and 2. Large differences (top ten most dissimilar feature weight) represent “distinct” functional connections, while small feature-weight differences (top ten most similar) represent “shared” network profiles. Both the distinct and shared network profiles are important when considering cognitive development as connections that change are equally important as connections that do not change between early and middle childhood.

**Figure 1:**
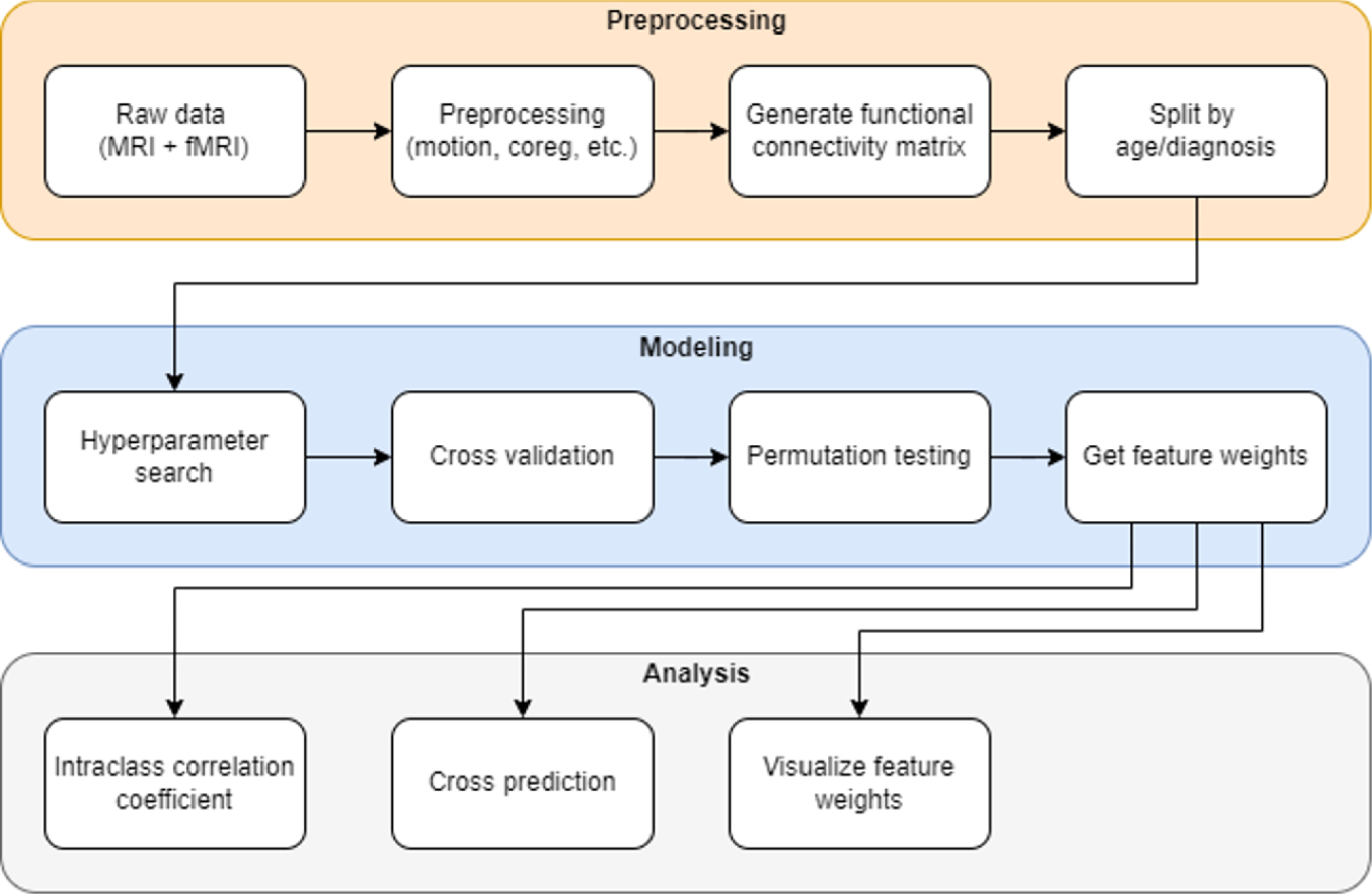
Processing stages for the neuroimaging data. There are three overall stages to the data pipeline: preprocessing, modeling, and analysis. Preprocessing involved correcting the raw MRI and fMRI data for motion, coregistering the structural and functional images, normalizing to a standard template, generating a functional connectivity matrix, and splitting the participants by age or diagnosis. Next is modelling and it starts with searching for the optimal parameters for the model, then training and validating the model using the functional connectivity matrices, randomly permutating the data, and ends with extracting the model’s feature weights. Lastly, the feature weights were analyzed by calculating the intraclass correlation coefficient, using the weights to cross-predict cognition in a different age bin, and visualizing the feature weights.

## Results

### Predicting age and sex in ADHD and NT

Using Ridge regression, we predicted the age (r2=0.45, p=.01) and sex of individuals in the ADHD group (n=373) with an accuracy of 74% (p<.001). Age (r2=0.13, p=.01) and sex (60% accuracy, p=.01) were also predicted in the NT group (n=106). We class-balanced the sex prediction to ensure that the model was not constantly predicting the most prevalent sex.

### Predicting cognitive ability in ADHD and NT

Using Ridge regression, we found the model could predict FSIQ (r=0.38, p=.002), VSI (r=0.31, p=.002), VCI (r=0.39, p=.002), FRI (r=0.30, p=.002), and WMI (r=0.21, p=.004), but failed to predict PSI (r=0.05, p=.26) in the group of participants diagnosed with ADHD (n=373). Conversely, we could not predict FSIQ (r=0.04, p=.42), VSI (r=0.16, p=.11), VCI (r=0.20, p=.05), FRI (r=-0.07, p=.73), WMI (r=0.12, p=.21), and PSI (r=-0.06, p=.70) in the NT group (n=106). These p-values were corrected for multiple comparisons using the max-statistic method. To determine whether these results were driven by model choice (Table 2), the analysis was replicated using a partial least squares (PLS) model. We found no difference in performance between the two models, except that VCI (r=0.23, p=.04) could be predicted in the NT group using the PLS model. The similarity between the two models was further supported by the ICC analysis which showed the weights produced by the Ridge and PLS model were strongly correlated (> 0.90; Supplement for additional details). Based on these results, we excluded the NT group, and used Ridge Regression for all subsequent analyses.

**Table 2:** Scores for predicting six cognitive abilities in ADHD and TD using partial least squares and Ridge regression. The Pearson r correlations test score represents the linear correlation between the model’s predicted values of the cognitive ability and the true values. The p-value was calculated by comparing the observed Pearson r score to a null distribution of Pearson r scores generated from 500 random permutations of the dataset. Both the PLS and Ridge models predicted FSIQ, VSI, VCI, FRI, and WMI in the ADHD group at significance (p<.011) but failed to predict PSI (p=.23). For the TD group, only VCI was predicted at significance (p=.04) using PLS. PLS and Ridge achieved similar Pearson r correlation scores on both the ADHD and TD groups. *values indicate statistically significant at p<.05 max-statistic corrected.

|  | ADHD (n=373) |  |  |  | TD (n=106) |  |  |  |
| --- | --- | --- | --- | --- | --- | --- | --- | --- |
|  | PLS |  | Ridge |  | PLS |  | Ridge |  |
| WISC Primary Index | Pearson r | P-value | Pearson r | P-value | Pearson r | P-value | Pearson r | P-value |
| Intelligence Quotient (FSIQ) | 0.37 | .002* | 0.38 | .002* | 0.04 | .388 | 0.04 | .417 |
| Visual Spatial (VSI) | 0.28 | .002* | 0.31 | .002* | 0.14 | .129 | 0.16 | .107 |
| Verbal Comprehension (VCI) | 0.37 | .002* | 0.39 | .002* | 0.23* | .035* | 0.20 | .051 |
| Fluid Reasoning (FRI) | 0.30 | .002* | 0.30 | .002* | -0.07 | .737 | -0.07 | .734 |
| Working Memory (WMI) | 0.17 | .011* | 0.21 | .004* | 0.08 | .259 | 0.12 | .213 |
| Processing Speed (PSI) | 0.06 | .227 | 0.05 | .257 | -0.10 | .791 | -0.06 | .698 |

Across all cognitive measures, models consistently assigned the largest positive weights to connections within two networks: memory retrieval and sensory/somatomotor (mouth), while inter-network connections with the largest positive weights were between memory retrieval and dorsal attention, and between memory retrieval and cerebellar. The largest negative weights were commonly assigned to connections between frontoparietal task control and visual, between dorsal and ventral attention, and to networks connected with subcortical areas.

### Developmental changes linking functional connectivity and cognitive abilities in ADHD

To examine developmental changes in the relationship between neural connectivity profiles and cognitive ability, we divided the ADHD group into three age bins (Table 3). The model successfully predicted FSIQ (r = 0.27, p = 0.02), VSI (r = 0.24, p =0.02), and VCI (r = 0.22, p = 0.03), but not FRI and WMI (p > 0.05) for Bin 1 (ages 6-8); and FSIQ (r = 0.35, p = 0.002), VSI (r = 0.21, p=.02), VCI (r = 0.35, p = 0.002), FRI (r = 0.31, p = 0.004), and WMI (r = 0.29, p = 0.004) for Bin 2 (ages 9-11). The model did not predict any WISC-V measure (p > 0.17) for individuals in Bin 3 (ages 12-16). We found similar results using a smaller sample of Bin 2 (n=113) that matched the sample sizes of Bins 1 and 3; the model could predict FSIQ (r = 0.37, p = 0.002), VSI (r = 0.27, p = 0.01), VCI (r = 0.37, p = 0.002), FRI (r = 0.30, p = 0.006), WMI (r = 0.35, p = 0.002), but not PSI (r = 0.04, p = 0.38). The feature weights are shown in Figure 2.

**Table 3:**
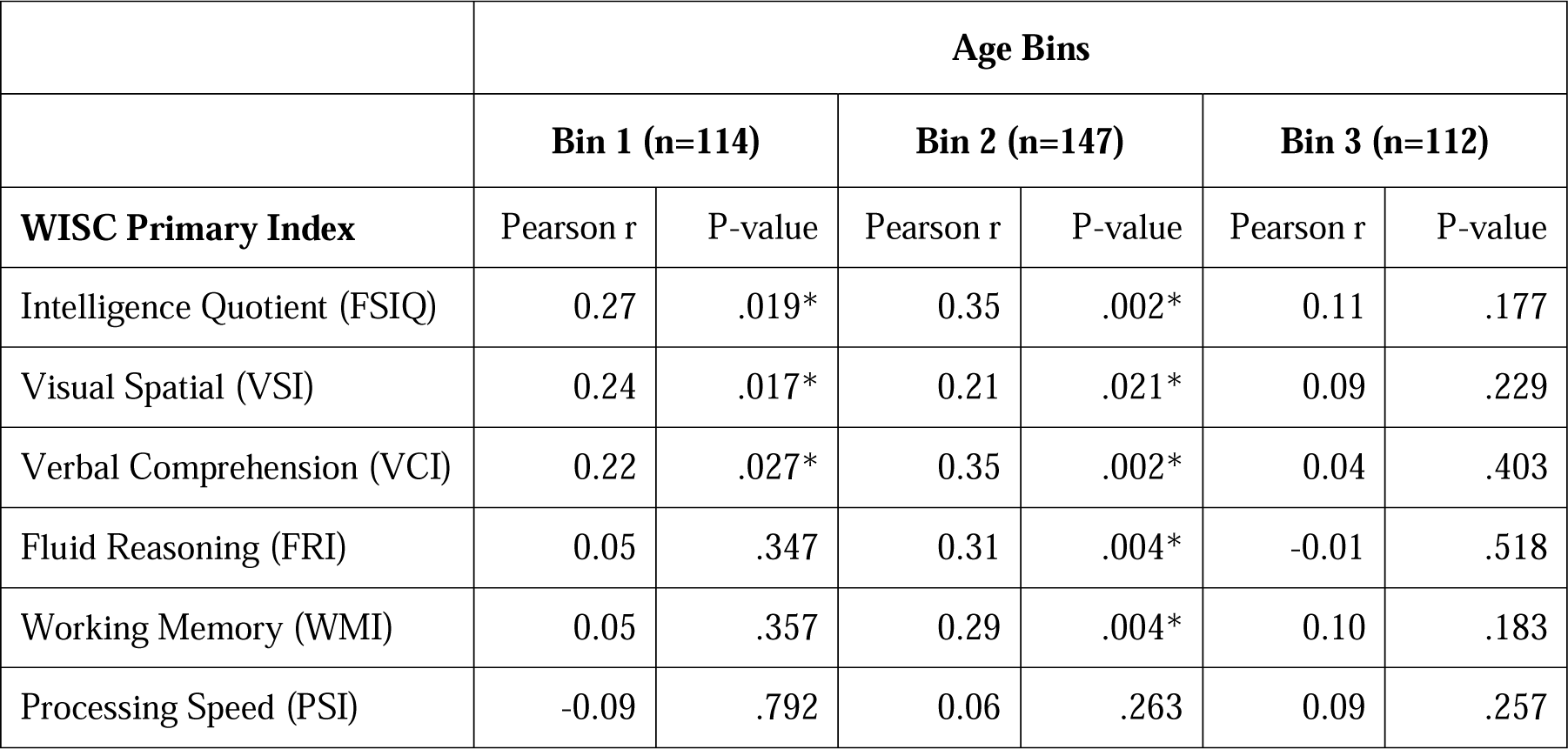
Scores for predicting six cognitive abilities in ADHD across three age bins using Ridge. Bin 1 represents early childhood (ages 6-8), Bin 2 represents middle childhood (ages 9-11), and Bin 3 represents adolescence (ages 12-16). The Ridge model successfully predicted FSIQ, VSI, and VCI in Bin 1 (p<.03); FSIQ, VSI, VCI, FRI, and WMI in Bin 2 (p<.02); and no cognitive ability in Bin 3 (p>.17). *values indicate statistically significant at p<.05 max-statistic corrected.

|  | Age Bins |  |  |  |  |  |
| --- | --- | --- | --- | --- | --- | --- |
|  | Bin 1 (n=114) |  | Bin 2 (n=147) |  | Bin 3 (n=112) |  |
| WISC Primary Index | Pearson r | P-value | Pearson r | P-value | Pearson r | P-value |
| Intelligence Quotient (FSIQ) | 0.27 | .019* | 0.35 | .002* | 0.11 | .177 |
| Visual Spatial (VSI) | 0.24 | .017* | 0.21 | .021* | 0.09 | .229 |
| Verbal Comprehension (VCI) | 0.22 | .027* | 0.35 | .002* | 0.04 | .403 |
| Fluid Reasoning (FRI) | 0.05 | .347 | 0.31 | .004* | -0.01 | .518 |
| Working Memory (WMI) | 0.05 | .357 | 0.29 | .004* | 0.10 | .183 |
| Processing Speed (PSI) | -0.09 | .792 | 0.06 | .263 | 0.09 | .257 |

**Figure 2:**
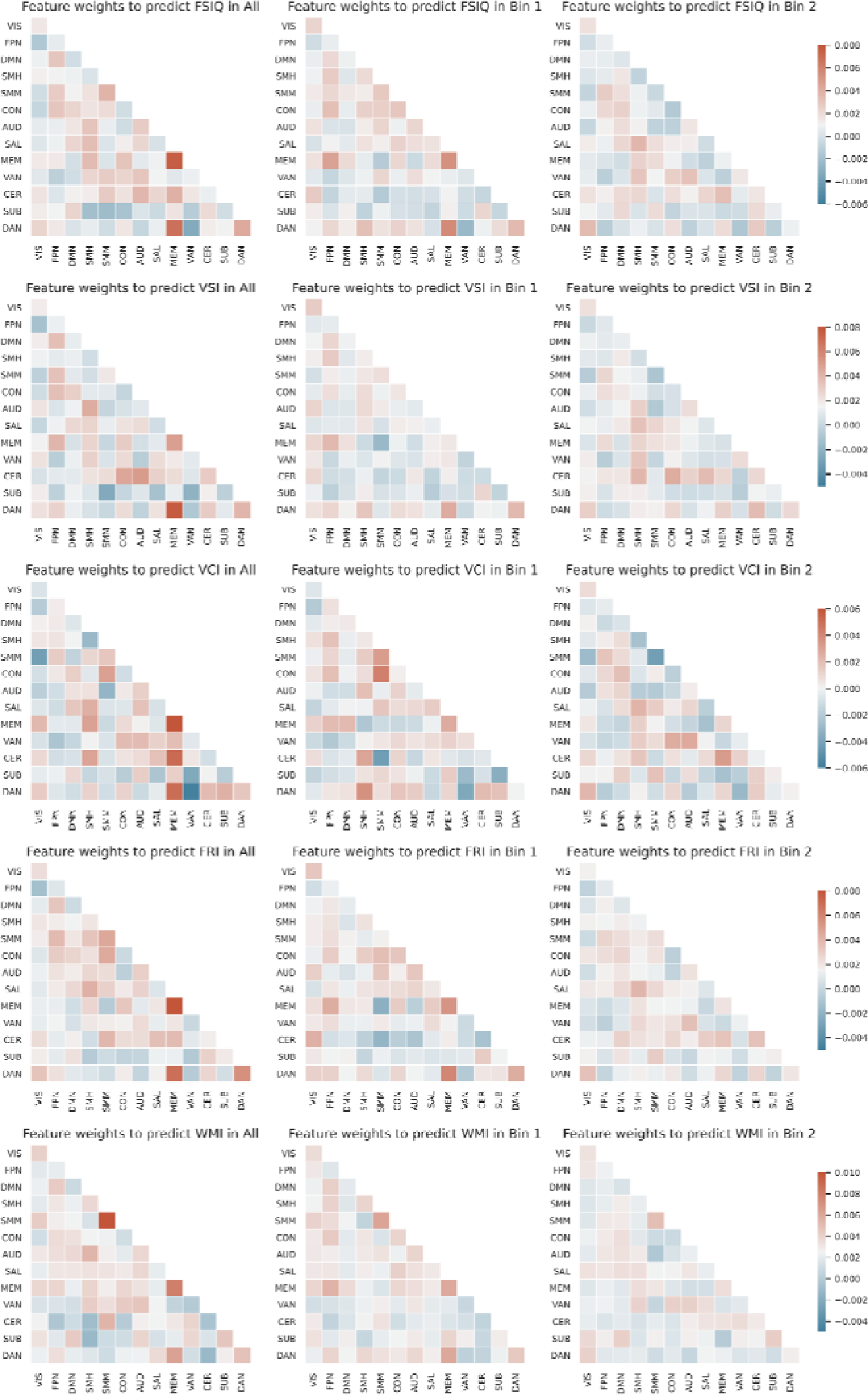
Feature weights used to predict five cognitive abilities in the entire ADHD group, Bin 1, and Bin 2. Each row represents one of five WISC measures: FSIQ, VSI, VCI, FRI, and WMI. Each column represents one of three ADHD groups: All (ages 6-16), Bin 1 (ages 6-8), and Bin 2 (ages 9-11). Each scale applies to the feature weight matrices in the row. A feature weight matrix represents the average feature weight for all connections between two networks shown for all networks. Darker cells in the feature weight matrix represent more extreme values, while lighter cells represent values closer to zero. Red cells represent positive values (increases in value for that network connection increased the predicted cognitive score), while blue cells represent negative values (increases in value for that network connection decreased the predicted cognitive score). Diagonal cells represent intranetwork connections, while off-diagonal cells represent internetwork connections. The networks are visual (VIS), frontoparietal task control (FPN), default mode (DMN), sensory/somatomotor (hand; SMH), sensory/somatomotor (mouth; SMM), cingulo-opercular task control (CON), auditory (AUD), salience (SAL), memory retrieval (MEM), ventral attention (VAN), cerebellar (CER), subcortical (SUB), and dorsal attention (DAN).

The feature weights for FSIQ, VSI, and VCI had positive weights for network connections between memory retrieval and dorsal attention, cingulo-opercular and sensory/somatomotor (mouth) networks, and within memory retrieval, sensory/somatomotor (mouth) networks. Negative weights were learned for connections between dorsal and ventral attention, memory retrieval and sensory/somatomotor (mouth), and between cerebellar and sensory/somatomotor (mouth) networks. For the Bin 2 feature weights, we found a general pattern of less-extreme feature weights (fewer darker-colored cells) across all cognitive measures (Figure 2, right column) relative to the entire ADHD group and Bin 1. This suggests that the model is not relying on specific network connections, but instead is using a distributed approach to predict cognitive ability. However, the model identified strong negative weights for connections within the sensory/somatomotor (mouth) network associated with predicting VCI scores, implying this network is deemphasized for predicting VCI performance. Interestingly, this connection was assigned a large positive in Bin 1, which shows that the models switched from a positive to a negative weight from Bin 1 to Bin 2 when predicting VCI (see Supplement for additional details).

### Cross-prediction across age bins in ADHD

Using cross-prediction (out-of-sample cross-validation), we found that models trained on Bin 1 and tested on Bin 2 successfully predicted FSIQ (r=0.33, p=.002), VSI (r=0.36, p=.002), VCI (r = 0.32, p = 0.002), and FRI (r = 0.15, p = 0.02) (Figure 3). We also found the reverse; a model trained on Bin 2 and tested on Bin 1 successfully predicted FSIQ (r = 0.36, p = 0.002), VSI (r = 0.40, p = 0.002), VCI (r = 0.30, p = 0.002), and FRI (r = 0.20, p = 0.01). However, models failed to cross-predict WMI (r = 0.03, p = 0.35) and PSI (r = 0.07, p = 0.18) when trained on Bin 1 and tested on Bin 2, and when trained on Bin 2 and tested on Bin 1; WMI (r = 0.03, p = 0.37) and PSI (r = 0.03, p = 0.40). These results suggest connectivity patterns associated with FSIQ, VSI, VCI, and FRI, but not WMI and PSI, generalize from early to middle to childhood.

**Figure 3:**
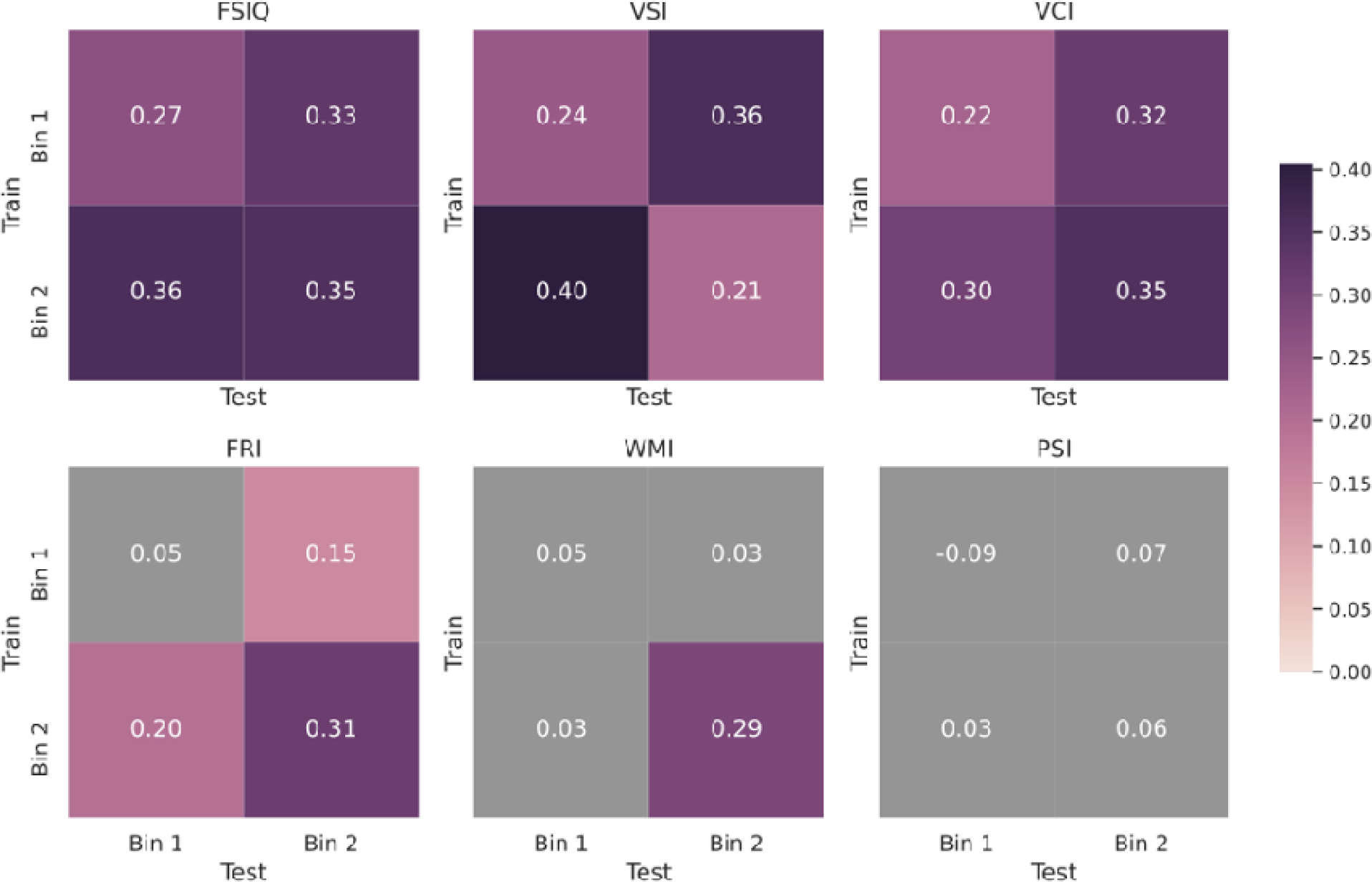
Scores for cross-predicting six cognitive ability between Bin 1 and Bin 2. For each matrix, rows represent the age bin (Bin 1 or Bin 2) the model was trained on, while columns represent the age bin (Bin 1 or Bin 2) the model was tested on. The top-left to bottom-right diagonal represents training and testing the model within the same age bin (same scores as in Table 3), while the bottom-left to top-right diagonal represents the training the model on Bin 2 and testing on Bin 1 and training the model on Bin 1 and testing on Bin 2 respectively. Values within each cell are the Pearson r correlation test score and represent the linear correlation between the model’s predicted values of the cognitive ability and the true values. Purple cells indicate statistically significant at p<.05 after being corrected for multiple comparisons using the max-statistic method, while grey cells indicate not statistically significant.

To identify the most similar and dissimilar feature weights that were trained on Bin 1 and Bin 2, we subtracted (using absolute values) the Bin 2 feature weights from the Bin 1 feature weights for each cognitive measure (e.g., for FSIQ, VSI, VCI, FRI, and WMI). We found the feature weight profiles with the (top ten) most similar networks between early childhood (Bin 1) and middle childhood (Bin 2) across all cognitive measures comprised of four intra-network connections: the frontoparietal, default mode, subcortical, and dorsal attention networks. The feature weights associated with inter-network connections that were most similar between the two age groups primarily included the frontoparietal, default mode, subcortical, and salience structures, although other networks were also found (but to a lesser degree) to be shared between age groups.

Conversely, relatively more intra-network connections were dissimilar between the two age groups across the cognitive measures, such as the sensory/somatomotor (mouth), cingulo-opercular, and memory retrieval networks, but also included cerebellar and ventral attention networks. Most dissimilar (top ten) inter-network connections included the memory retrieval, dorsal attention, sensory/somatomotor networks (mouth and hand) networks. Moreover, connections between cingulo-opercular network and other parts of the brain were more often shared than not between the age groups across the different cognitive measures. Note, the model was not able to predict FRI and WMI in Bin 1 but was able to predict FRI and WMI in Bin 2, which does not reflect direct comparisons of specific cognitive abilities between the age groups (Figure 4).

**Figure 4:**
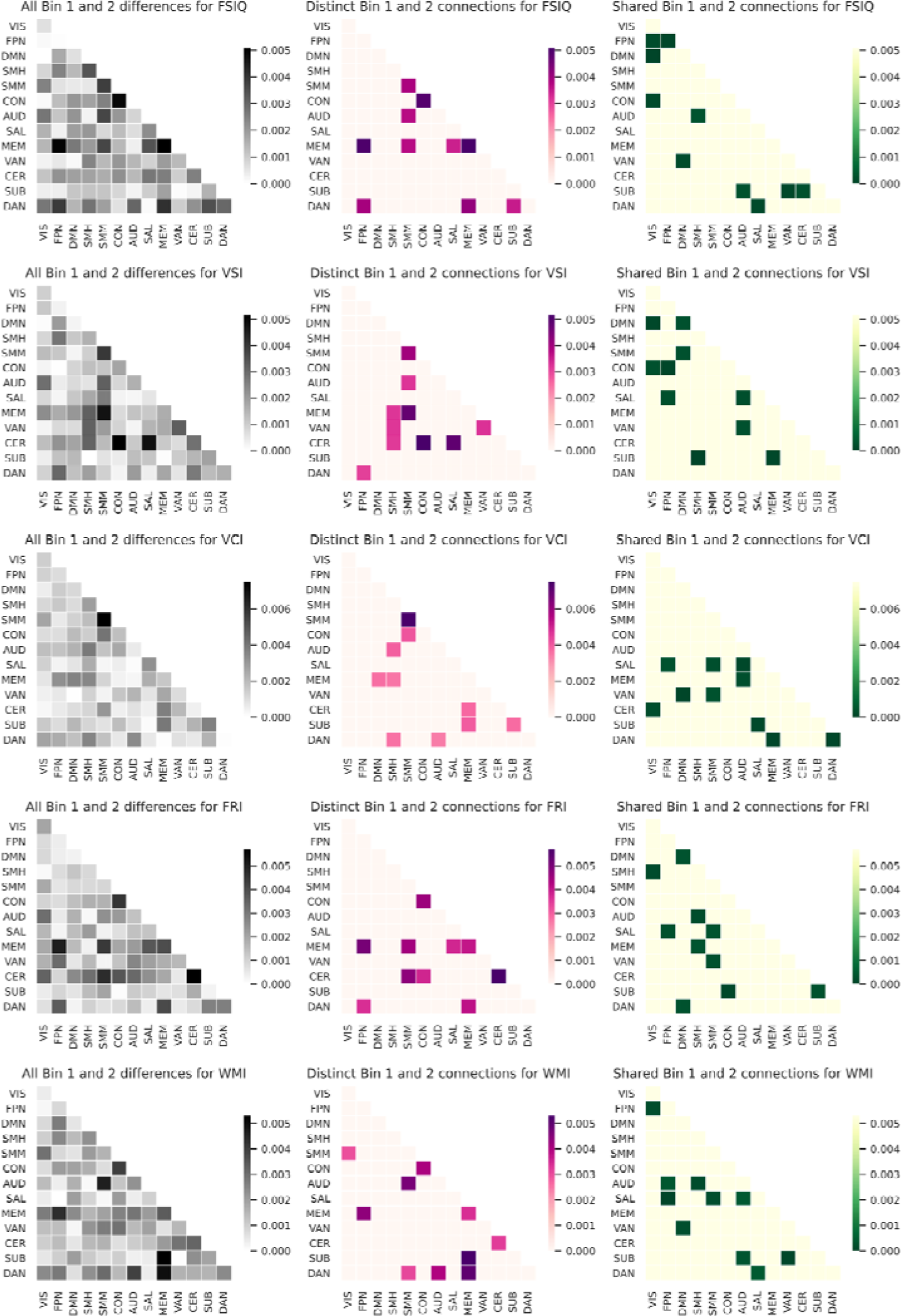
Difference in feature weights between Bin 1 and Bin 2 for five cognitive abilities. Each row represents one of five WISC measures: FSIQ, VSI, VCI, FRI, and WMI. The left column (grey) represents all feature weight differences between Bin 1 and 2, the center column (pink) represents network connections with the most dissimilar values for Bin 1 and 2 (“distinct” networks), and the right column (green) represents network connections with the most similar values between Bin 1 and 2 (“shared” networks). The distinct network profiles were obtained by thresholding all feature weight differences between Bins 1 and 2 by the ten largest differences. The shared network profiles were obtained by thresholding all feature weight differences between Bins 1 and 2 by the ten smallest differences. For the left and center columns, darker cells represent a larger difference between the feature weights assigned to Bin 1 and 2 when predicting cognition, while lighter cells represent a smaller difference. Diagonal cells represent intra-network connections, while off-diagonal cells represent internetwork connections.

## Discussion

We demonstrated that applying machine learning to movie-watching fMRI data is a viable tool for predicting demographic and higher-level cognitive abilities in children and adolescents diagnosed with ADHD. In a large cohort of early childhood, middle childhood, and adolescent participants, we built models that successfully predicted age and sex, and we identified shared and distinct neural mechanisms associated with different aspects of higher-level cognition across development in ADHD and NT groups.

Establishing models that can predict age and sex is important because it demonstrates that a dimensional data-driven approach (i.e., machine learning) can be used to extract information from the neural connectivity profile to predict aspects of development. The distinct feature-weight profile used by the model likely reflects that male and female (sex was limited to these two categories) children and adolescents have distinct functional patterns of brain activity and are relying on different neural mechanisms to process the movie. This result replicates and expands on previous work that generated models to predict age and sex^25,42–44^ providing further evidence that biological properties, such as age and sex, could be reliably localized to specific important connections in the brain. Does the same apply to cognition?

Emerging from our results was predicting higher-level cognitive abilities followed an inverted-U pattern. This suggests that the link between intra brain functional connectivity in response to movie watching and cognitive ability is strongest and most consistent during middle childhood, weaker and more variable during early childhood and not detectable during adolescence. Although successfully predicting cognitive abilities based on neuroimaging data is consistent with previous studies (in neurotypical children^45^), we were surprised that the same cognitive systems could not be predicted in adolescence with ADHD. One reason for this pattern of results is that the link between functional connectivity and cognitive ability in adolescence may be too weakly represented in the data. This could reflect more variable cognitive development in this group that was not equally elicited by the movie. Another interesting possibility is that the nature of the relationship between cognition and connectivity profiles do not follow a linear trajectory and linear models are insufficient to capture the link.

Although we could generate models that predicted different cognitive abilities in early and middle childhood, not all cognitive systems were reliably predicted between the cohorts. Beyond that, the models that best predicted cognitive abilities were different for participants in early versus middle childhood. For example, the model could predict fluid reasoning and working memory in middle childhood but not in early childhood. Although previous work has suggested that verbal and visuospatial working memory remain relatively distinct in children ages 4 to 11^46^, our results suggest this may not be the case. One potential reason why we could predict working memory in middle, but not early childhood, is because the link between these cognitive abilities and the underlying neural mechanisms are either weaker or follow distinct developmental trajectories during this period in children with ADHD^7,47^. Indeed, maturing working memory and fluid reasoning are associated with the development of the frontoparietal network^48,49^, and atypical development of this network is associated with difficulties in fluid reasoning and working memory in children between the ages of 6 and 12 with ADHD^50–52^.

One similarity across the three developmental stages was that processing speed could not be predicted well in children and adolescents with ADHD. This is likely due to children with ADHD showing the most pronounced deficits in processing speed^7,47^. Delayed or more variable development of this cognitive ability may suggest that there is a weakened relationship between neural activity associated processing speed ability, or the representation of processing speed in brain activity may have greater variability, resulting in poorer prediction scores. Similarly, previous work found the lowest prediction scores on measures of speed/flexibility out of three higher-order cognitive functions (General Ability, Speed/Flexibility, and Learning/Memory) using resting-state fMRI data in 9-to 10-year-old children^45^. These results suggest that neither movie-watching or resting-state fMRI is ideal for capturing the neural mechanisms related to processing speed in neurotypical children^53^ or children with ADHD.

We hypothesized models identifying unique network patterns associated with cognitive ability in one age cohort would not generalize to other cohorts. However, this was not the case. Instead, connectivity patterns associated with IQ, visual spatial, verbal comprehension, and fluid reasoning generalize from early to middle to childhood. The shared networks for predicting cognition between early and middle childhood were divided into two types: intra-network (within) connections and inter-network (between) connections. Shared intra-network connections were predominately made up of the frontoparietal, default mode, subcortical, and dorsal attention, but also included sensory (e.g., visual and auditory) networks. The shared inter-network connections were comprised primarily of the frontoparietal, default mode, memory retrieval, dorsal attention, and salience networks. This is not to say these network connections remain stable during this period; some developmental changes may not be tied to cognitive ability. What the shared network connections do imply is that the models did not change their importance for these network connections when predicting cognition for early and middle childhood. Why would the models highlight these networks? One possibility is that many of the shared networks, which have been linked to higher-level cognitive processing— such as the frontoparietal, memory retrieval, dorsal attention, and salience networks—bridge cognitive maturity and the degree to which they are recruited during movie watching is similar between the two age groups. That is, young children with greater scores on cognitive abilities are recruiting (or not recruiting) these networks during movie watching to the same degree as older children, while the same relationship is true for early and middle childhood participants with lower scores on cognitive abilities.

Although we found many important shared network connections between the two age groups, relatively low explained variance suggests that the shared networks do not capture all, or even most, of the developmental neural mechanisms supporting higher-level cognition in early and middle childhood. The distinct networks for predicting cognition between early and middle childhood were found primarily within the sensory/somatomotor (mouth), cingulo-opercular, and memory retrieval networks, but also included subcortical, cerebellar, and ventral attention networks, and between the sensory/somatomotor networks (mouth and hand), dorsal attention, and memory retrieval networks. In line with our results, the sensory/somatomotor (hand) and memory retrieval networks connections were important when predicting general cognitive ability in a cohort of middle childhood participants^45^. This suggests the network connectivity profile of the sensory/somatomotor (hand), and memory retrieval networks are strongly linked to cognitive development from early to middle childhood. Contrary to our hypothesized outcomes, the connectivity profile in frontoparietal network was not distinctly associated with cognition in early versus middle childhood. Although the frontoparietal network is strongly linked to the development of executive function and intelligence^54,55^, it’s possible that Despicable Me did not trigger the cognitive systems mediated by the frontoparietal network in the youngest two cohorts in our study. Another possibility is that movie-watching in general may not be a context that is sensitive enough to extract neural features associated with executive functioning in young children. Related to both points, previous studies found that the frontoparietal network is not a flexible hub during movie watching^27^ to the same degree it has been reported to be when participants complete a set of demanding tasks^56,57^. Therefore, the bridge between frontoparietal activity associated with executive functioning during movie-watching and the WISC scales may not be sufficiently strong to find patterns of activity associated with executive functioning development.

Importantly, we were able to replicate all findings using a different model: partial least squares. In fact, we found a very high correspondence between the feature weights generated by Ridge and partial least squares as measured by the high intraclass correlations. This suggests that the models’ ability to predict cognition is likely not driven by model choice as both the model output (its correlation score) and the model internals (its feature weights) are extremely similar between Ridge and partial least squares. In line with our results, previous studies also reported finding little difference between a Lasso model and Ridge’s correlation score when predicting fluid and crystalized intelligence^25^. Thus, perhaps in the space of regularized linear models, the choice of model does not lead to significant performance differences.

### Limitations and Future Directions

One limitation of the current study is that we were unable to predict cognition in age-matched neurotypical children, despite previous studies demonstrating that cognition can be predicted in this populations^25,29,45^. One potential reason is the NT group was smaller than the ADHD group. However, this likely does not account for our results because we were able to predict some cognitive measures in the ADHD group with a comparable sample. Another factor might be data quality; perhaps the noisier data for the NT group was leading to poor predictive performance. This is also an unlikely to account for our results because we could predict age and sex in the NT group. However, the explained variance and accuracy was lower in the NT group compared to the ADHD group, and lower than estimates from other studies (approximately 42% explained variance but using a different task^44^). To determine whether our findings are specific to ADHD or generalize to other groups of children, future studies examining distinct neural mechanisms associated with cognitive development in neurotypical populations should replicate our findings using larger samples. Furthermore, different models would be valuable, such as those incorporating non-linear relationships between connectivity profiles and cognition, and lesion-modeling^58^ that examine changes in the direction of the relationship between neural mechanisms and cognition across development.

### Conclusion

Different higher-order cognitive abilities in a large group of children and adolescents diagnosed with ADHD could be predicted using functional neural activity during movie watching. Prediction scores do not remain constant across development but instead follows an inverted-U developmental trajectory from early childhood to adolescence, and that certain neural mechanisms linked to higher-level cognition were shared we also found several distinct sets of neural mechanisms for predicting cognition between early and middle childhood.

## Supporting information

Suppmentary Material

## Acknowledgements

We would like to thank the Child and Mind Institute for designing and collecting data for the Healthy Brain Network Biobank. We would also like to thank the children, adolescents, and their families for taking the time to participate in studies conducted by the Child and Mind Institute. This work was funded by a Natural Sciences and Engineering Research Council of Canada Discovery grant (RGPIN-2020-05042 to BS), and a Vector Institute Research Grant, and BrainsCAN research grant to (YM). The authors report no biomedical financial interests or potential conflicts of interest.

