## Supplementary material for "Identifying Developmental Changes in Functional Brain Connectivity Associated with Cognitive Functioning in Children and Adolescents with ADHD": Suppmentary Material

**Supplement**

**Data Pipeline**

An overview of the pipeline for the neuroimaging data is shown in Supplement Figure 1. There were three overall stages in the pipeline: preprocessing, modeling, and analysis. The preprocessing stage was used to clean the data of artifacts and noise, and to generate functional connectivity matrices. The modeling stage was when the data was used to train machine learning models to predict cognitive ability from functional connectivity. The last stage, analysis, extracted the model's feature weights to gain insights into the neural mechanisms underlying cognition and cognitive development. For the phenotypic data, we only excluded participants with an incomplete WISC assessment or with an FSIQ below 70.

**Participants**

Healthy Brain Network (HBN) biobank (releases 1 to 8), which is a part of the Child Mind Institute (Alexander et al., 2017), an ongoing initiative to create and share multimodal data from thousands of New York City children and adolescents between the ages of 5 to 21. The biobank uses a community-referred recruitment model with advertisements to parents, community members, educations, and local care providers. Participants were screened and excluded if there were impairments that would interfere with the study procedure, safety concerns, or medical concerns.

A total of 1,116 participants were initially downloaded from the HBN. Each data set was run through the initial preprocessing stages, and 880 of them met quality assurance thresholds, and functional connectivity matrices were generated for each data set. Subsequently, all structural and functional MRI data, including the functional connectivity matrices, were visually inspected for artifacts. Of the 880, 154 participants were excluded for poor data quality, leaving 726 datasets. An additional 18 participants were also excluded for having an FSIQ below 70. The last step was to filter for diagnosis: In total, 229 participants had a diagnosis that was not ADHD, 373 participants were diagnosed with ADHD (including comorbidities), and 106 participants had no clinical diagnosis (neurotypical (NT) group).

*Age selection*

Group assignment was based on rounding down the age provided by the HBN phenotypic profile. For example, a child with an age of 8.9 years old was categorized as an 8-year-old and grouped into Bin 1. The age 9 cutoff between Bins 1 and 2 was chosen because around this age is when children enter school and when symptoms of neurodevelopmental disorders not previously noticed begin to appear (Berger, 2017). The age 12 cutoff between Bins 2 and 3 is important because the diagnosis of ADHD in the DSM-V states that symptoms must start before the age of 12 and this age is typically when puberty begins, marking the start of adolescence (American Psychiatric Association, 2013; Berger, 2017).

*MRI acquisition and preprocessing*

All MRI data was collected on a 3 T Siemens scanner using a Siemens 32-channel head coil. The structural MRI scans were acquired in 224 sagittal (TR=2500 ms, TE=3.15 ms, resolution=0.8 x 0.8 mm2). The functional MRI scans were acquired with a gradient-echo planar imaging pulse sequence (TR=800 ms, TE=30 ms, Flip Angle=31 degrees, whole brain coverage 60 slices, resolution=2.4 x 2.4 mm2). Preprocessing of functional data included motion correction (using six motion parameters: left/right, anterior/posterior, superior/inferior, chin up/down, top of head left/right, nose left/right), and functional and structural scans were co-registered and normalized to the Montreal Neurological Institute (MNI) template. Functional data were then spatially smoothed using a Gaussian filter (8 mm kernel) and low-frequency noise, such as drift, was removed by high-pass filtering with a threshold of 1/128 Hz. The data was denoised using Bandpass filter regressors with cerebrospinal fluid, white matter signals, motion parameters, their lag-3 2nd-order volterra expansion (Friston et al., 2000), and spikes (based on mean signal variance across volumes) as nuisance regressors.

*Cognitive abilities*

Five cognitive abilities were collected from the Wechsler Intelligence Scale for Children Fifth Edition (Wechsler, 2014). Visual Spatial Index (VSI) measures the ability to evaluate visual details and understand visual spatial relationships to construct geometric designs from a model. Verbal Comprehension Index (VCI) measures the ability to access and apply acquired word knowledge. Fluid Reasoning Index (FRI) measures the ability to detect the underlying conceptual relationships among visual objects and requires reasoning to identify and apply rules. Working Memory Index (WMI) measures the ability to register, maintain, and manipulate visual and auditory information in conscious awareness. Processing Speed Index (PSI) measures the speed and accuracy of visual identification, decision making, and decision implementation. Full-Scale IQ (FSIQ) represents general intellectual ability across a diverse set of cognitive abilities. Both the five primary indices and FSIQ are on a standard score metric with a mean of 100 and a standard deviation of 15. The primary index scores range from 45 to 155, while the FSIQ score ranges from 40 to 160. For both the primary indices and FSIQ, scores between 90 to 109 are considered average for a typically developing child.

*Partial least squares*

Partial least squares (PLS) is a statistical method of finding a linear regression model by projecting the samples (X) and the targets (y) to a new latent space. A PLS model will search the multidimensional sample space that explains the maximum multidimensional variance in the target space. That is, PLS searches for the latent variables most strongly associated between the sample and target data. For this study, I used the univariate version of PLS, which is a form of regularized linear regression. The univariate version is in the same class as Ridge regression and principal component regression where the number of components controls the strength of regularization (Wegelin, 2000).

*Ridge regression*

Ridge is another method in the class of regularized linear regression but instead of using partial least squares, it uses complete least squares with L2 regularization. Regularization is a technique to impose constraints on the feature weights of a linear model to reduce overfitting on the data, thereby improving generalizability. L2 regularization, also known as Ridge, implements regularization by adding the square of the absolute value of the feature weights in the least-squares penalty. This penalty on the size of feature weights enables the model to be more robust to the collinearity of features by making the feature weights sparser.

*Hyperparameter Search*

A hyperparameter search was performed to determine the optimal number of components (between two and six) for the PLS model. We found four components resulted in the highest test score on all cognitive measures. To determine the optimal alpha value for the Ridge model, a hyperparameter search was done by varying alpha and selecting the value that resulted in the highest test score. The explored range for alpha was between 1 to 10,000 (exclusive) with a step size of 100. The hyperparameter search was conducted using the 10-fold 10-repeat cross validation scheme described in the Supplement. The success of the model to predictive cognitive ability based on functional connectivity profiles was evaluated using permutation statistics using the max-statistic method (Nichols & Hayasaka, 2003; see Supplement for more details) on the Pearson correlation (Finn & Bandettini, 2021, Sripada et al., 2020, and Tian & Zalesky, 2021).

*Cross validation*

After optimizing for the component parameter for PLS and the alpha parameter for Ridge, both models were cross-validated using a 10-fold 10-repeat scheme. The scheme starts by splitting the dataset into 10 equal chunks where 9 of the chunks are used to train the model and the remaining 1 chunk is used to test the model. This process is repeated 9 times where each chunk is used for testing and the rest are used for training, resulting in 10 folds and 10 Pearson r scores. After these 10 folds, the dataset is shuffled and a new set of 10 folds is generated. This process is repeated 9 more times, resulting in 100 variations in the training and testing set and 100 Pearson r scores. The final cross-validation score is the mean of these 100 Pearson r scores. Cross-validation was performed to avoid overfitting as the model's performance on a random sampling of the data may not represent the model's predictive performance on unseen data. Thus, by splitting the dataset into 100 variations for training and testing, I achieved a more robust and accurate estimate of the model's performance on the data.

*Permutation testing*

Permutation statistics were used to evaluate the significance of the cross-validation score, which is the correlation between the predicted and actual cognitive scores. A permutation-based p-value represents how likely the performance of a model— the cross-validation score—would be obtained by chance. The null hypothesis is that the model fails to find any statistical dependency between features (functional connectivity) and targets (cognitive ability). This manifests as the model predicts random cognitive scores, which is reflected in random correlations between the predicted and actual cognitive scores. A null distribution was generated (reflecting the null hypothesis) by randomly permuting the targets (breaking dependencies between features and targets), and then training and testing the model on the shuffled dataset. Targets were shuffled 500 times, generating 500 permutation scores to form a null distribution. The p-value is calculated as the fraction of permutation scores greater than the cross-validation score. For example, if 5 of the 500 permutation scores were greater than the cross-validation score, then the calculation outputs a p-value of 0.01. A low p-value implies a low probability that the performance of the model was obtained by chance. A high p-value implies either a lack of dependency between features and targets, or that the model was not able to learn the dependency in the data. In this case, using a different model that can capture the dependency in the data may result in a lower p-value.

To correct for multiple hypotheses, we used the max-statistic method (Nichols & Hayasaka, 2003). This method combines the null distribution among a group of tests into one null distribution for the entire group. For this study, I combined the null distributions for all cognitive measures within each age group for each model. The 95th percentile value was used as the threshold for significance representing an alpha of 0.05. Cross-validation scores above the threshold were considered significant with max-statistic correction, while scores below the threshold were not considered significant.

*Intraclass correlation coefficient in ADHD*

The ICC is a measure of how correlated the same variables are in different observations and can range from zero to one with an ICC of one indicating identical values for variables in different observations. For our purposes, ICC was used to measure how similar the PLS feature weights were to the Ridge feature weights. Thus, we could evaluate whether both models were using similar or different feature weights to predict cognitive ability. This use-case of ICC is identical to a one-way ANOVA fixed effects model.

To explore the similarity between PLS and Ridge, we calculated the intraclass correlation coefficient (ICC) between each model's feature weights trained on the ADHD group for each cognitive ability (Table 4). The ICC revealed strong correlations (> 0.90) in the weights produced by the Ridge and PLS models, which suggest convergence between the features that PLS and Ridge find important. Specifically, the lowest ICC (0.92, p<.001) was found for Bin 1 in PSI, while the second-highest ICC (0.98, p<.001) was found for Bin 2 in FSIQ, WMI, PSI, and for Bin 3 in PSI. The highest ICC (0.99, p<.001) was found for Bin 1 on FRI. Thus, the consistency between model output (Table 2) and feature weights (Table 4) strongly suggests that both PLS and Ridge used near-identical network patterns to predict cognition.

| WISC Primary Index | All Ages | Bin 1 | Bin 2 | Bin 3 |
|---|---|---|---|---|
| Intelligence Quotient (FSIQ) | 0.96 | 0.97 | 0.98 | 0.97 |
| Visual Spatial (VSI) | 0.96 | 0.95 | 0.94 | 0.93 |
| Verbal Comprehension (VCI) | 0.96 | 0.97 | 0.96 | 0.96 |
| Fluid Reasoning (FRI) | 0.96 | 0.99 | 0.96 | 0.97 |
| Working Memory (WMI) | 0.95 | 0.96 | 0.98 | 0.93 |
| Processing Speed (PSI) | 0.96 | 0.92 | 0.98 | 0.98 |

**Supplementary Table 1: Intraclass correlation coefficients between partial least squares (PLS) and Ridge feature weights.**

The ICC describes how strongly values for the same variables resemble each other. A high ICC implies similar feature weights between PLS and Ridge, while a low ICC implies dissimilar feature weights. The lowest ICC was 0.92 (p<.001) when predicting PSI in Bin 1 and the highest ICC was 0.99 (p<.001) when predicting FRI in Bin 1. Overall, the high ICC values for all cognitive measures across all groups suggest that PLS and Ridge used very similar feature weights when predicting cognition. All values have a p<.001 and were corrected for multiple comparisons using FDR.

*Model Feature Weight Analysis and Cross-prediction*

Cross-prediction was performed by training a model on one age bin and testing it on another age bin. For instance, a model trained on Bin 1 with FSIQ was used to predict FSIQ in Bin 2. This was, the generalizability of the models could be examined, including shared and distinct feature weights across age bins. To perform the cross prediction, data sets in each of the age bins were split into training and testing sets using the 10-fold 10-repeat cross validation scheme. A model was trained on the in-group age bin and tested it on the out-group age bin. This process was repeated for all 100 folds and the final score was the average score from these folds. Permutation testing was also applied to this final score by randomly permuting the cognitive abilities within their respective groups and rerunning the modeling.

Model feature weights were extracted and visualized to better understand which network connections it found important to predict cognition. To obtain the final feature weights from the 10-fold 10-repeat cross-validation scheme, I averaged the 100 feature weights generated by each model in each fold. This resulted in a 34,716 x 1 feature weight vector where each value is the weight assigned to a feature. This node-level feature weight vector was condensed to a network-level feature weight vector by taking the mean of all connections between two networks to get one weight per network-network pair instead of one weight per node-node pair. By repeating this procedure for all 13 Power atlas networks, I ended up with a 13 x 13 network-level feature weight matrix. This reduction was performed to make the feature weights more interpretable and to focus on a network-level analysis. While these feature weights have information for understanding how the model predicts cognition, they do not have information on how cognition develops.

### Results

*Predicting cognitive ability in ADHD and NT*

For FSIQ, we found the model assigned large positive weights to connections between memory retrieval and dorsal attention networks, and connections within memory retrieval network. In contrast, the same model assigned large negative weights to connections between dorsal and ventral attention, subcortical connections with sensory/somatomotor (mouth and hand) and cingulo-opercular task control. For VSI, strong positive weights were assigned to connections between memory retrieval and dorsal attention network, and strong negative weights were assigned to connections between subcortical and sensory/somatomotor (mouth). For VCI, networks connected with the memory retrieval network, such as the cerebellar network positively predicted VCI performance, while connections between dorsal and ventral attention, ventral attention and subcortical, and visual with sensory/somatomotor (mouth) networks negatively predicted VCI performance. For FRI, we found a similar pattern of large positive weights as seen in FSIQ, namely connections between memory retrieval and dorsal attention networks, and connections within memory retrieval. A strong negative weight was found between frontoparietal task control and visual network. Lastly, WMI was best predicted by large positive weight for connections between the sensory/somatomotor (mouth) and in memory retrieval networks (Figure 2; left column).

*Developmental changes in links between functional connectivity and cognitive abilities in ADHD*

For the youngest participants (Bin 1), the model found that strong connections between memory retrieval and dorsal attention network, and connections within the memory retrieval network, were important positive predictors of FSIQ, while connections between dorsal and ventral attention were important negative predictors. In comparison, predicting VSI shared the same positive importance of the memory retrieval and dorsal attention connections as FSIQ, but the model differed in the strong negative weight for the connection between memory retrieval and sensory/somatomotor (mouth) network. VCI was best predicted based on strong positive weights for connections between cingulo-opercular and sensory/somatomotor (mouth) networks, and for within sensory/somatomotor (mouth) connections. Strong negative weights were learned for cerebellar and sensory/somatomotor (mouth), and for dorsal and ventral attention networks.

*Cross-prediction across age bins in ADHD*

The most distinct connectivity profiles associated with FSIQ were connections within the memory retrieval, cingulo-opercular, and sensory/somatomotor (mouth) networks and connections between the memory retrieval and frontoparietal, sensory/somatomotor (mouth), salience, and dorsal attention networks, in addition to connections between the dorsal attention network with the frontoparietal network. The most similar network profiles between the age groups included connections within the frontoparietal network, and connections between the ventral attention and default mode networks, between dorsal attention and salience networks, and between the visual network with the frontoparietal, default mode, and cingulo-opercular networks. We also found connections between subcortical areas and auditory, ventral attention, and cerebellum were similar between Bin 1 and 2.

The pattern of connections associated with VSI performance that were most dissimilar between the two age groups included connections within sensory/somatomotor (mouth), connections between sensory/somatomotor (mouth) and auditory networks, and memory retrieval, in addition to connections between dorsal attention and frontoparietal networks, and cerebellar connections to sensory/somatomotor (hand), cingulo-opercular, and salience networks. The feature weights that remained stable between the two age groups included connections with the default mode, cingulo-opercular, and frontoparietal networks.

The connectivity pattern linked to VCI scores that are most dissimilar feature weights between early childhood (Bin 1) and middle childhood (Bin 2) include connections within the sensory/somatomotor (mouth) and subcortical networks, and large differences in the feature weights were associated with connections between the sensory/somatomotor (hand) and the memory retrieval, the default mode, and the dorsal attention networks. Conversely, the most similar set of feature weights were connections within salience and dorsal attention internetwork connections, and connections between memory retrieval and auditory connections, as well as ventral attention network connections to the default mode network, and dorsal attention network connections to the memory retrieval network.

In contrast, dissimilar feature weights between the two age groups associated with predicting FRI included within cingulo-opercular, memory retrieval, and cerebellar network connections, while between-network connections included memory retrieval to frontoparietal, salience, and dorsal attention networks, in addition to connections between the dorsal attention network to the frontoparietal network. Interestingly, there were also large differences in feature weights with cerebellar connections, specifically to sensory/somatomotor (mouth) and cingulo-opercular networks. Key similar network configurations based on feature weights between the two age groups include within default mode and subcortical networks, and similar between networks include default mode network connections with the dorsal attention network, frontoparietal network connections with the salience network, and sensory/somatomotor networks (hand and mouth) connections to auditory, salience, memory, ventral attention, and visual attention networks.

Finally, the most dissimilar feature weights associated with predicting WMI include connections within the cingulo-opercular, memory retrieval, and cerebellar networks, which are the same cortical networks that were most dissimilar for predicting FRI. We also found dissimilar internetwork connections, notably between the dorsal attention network and the sensory/somatomotor (mouth), auditory, and memory retrieval networks, as well as between the frontoparietal network and the memory retrieval network. Feature weight profiles that were the most similar between the two age groups included connections between the salience network with the frontoparietal, sensory/somatomotor (mouth), auditory, and dorsal attention networks; connections between ventral attention network with the default mode network; and connections between the frontoparietal network with both auditory and visual networks. Interestingly, we found no within-network connections that were shared between the two age groups. Feature weights associated with the memory retrieval network were not shared between early and middle childhood.
